# Climate Surveys of Biomedical PhD Students and Training Faculty Members in the Time of Covid

**DOI:** 10.1101/2022.01.06.475246

**Authors:** Deepti Ramadoss, Meghan McCord, John P. Horn

**Affiliations:** Office of Graduate Studies

## Abstract

In July 2020, four months into the disruption of normal life caused by the Covid-19 pandemic, we assessed the institutional climate within the School of Medicine. Voluntary surveys were completed by 135 graduate students in 11 PhD-granting programs and by 83 members of the graduate training faculty. Several themes emerged. PhD students work hard, but the number of hours spent on research-related activities has declined during the pandemic. The students are worried about the pandemic’s impact on their research productivity, consequent delays in their graduation, and diminished future job prospects. Many late stage PhD students feel they do not have adequate time or resources to plan for their future careers. Symptoms of anxiety and/or depression are prevalent in 51% of the students, based on answers to standardized questions. Most students report they have strong mentoring relationships with their faculty advisors and like their programs, but they identify to a lesser extent with the medical school as a whole. Faculty think highly of their graduate students and are also worried about the pandemic’s impact upon productivity and the welfare of students. Students are interested in access to an Ombuds office, which is currently being organized by the medical school. Moving forward, the school needs to address issues of bias, faculty diversity, support for mentor training, professional development, and the imposter syndrome. We must also work to create a climate in which many more graduate students feel that they are valued members of the academic medicine community.

## Background and Goals

A national effort to reimagine the scope of PhD training in the biomedical sciences has emerged over the past ten years. Important new points of emphasis in this effort include mentoring and mentor training, support for career exploration and planning by trainees, and the development of strategies to improve the long-term wellness and resilience of trainees. Implementation of these activities has advanced through partnerships between the graduate training community and the NIH [1–3], NSF, AAMC GREAT group (https://www.aamc.org/professional-development/affinity-groups/great), professional scientific societies, foundations (e.g. Burroughs Wellcome Fund, JED Foundation, Howard Hughes Medical Institute), two national FOBGAPT meetings [4, 5], the Council of Graduate Schools (https://cgsnet.org/graduate-student-mental-health-and-well-being), and the National Research Mentoring Network [6, 7].

Mentoring, career planning and resilience are important because they all influence the learning environment. As a consequence, these factors may contribute to the overall climate of an institution by enhancing the health, happiness, scholarly productivity and overall satisfaction of trainees and their faculty mentors. This report presents data from parallel climate surveys completed in July 2020 by PhD students and by members of the graduate training faculty at the University of Pittsburgh School of Medicine. Simply stated, we sought to understand characteristics of our community that would serve as a baseline to help focus and evaluate future data-driven efforts to strengthen the local training environment for biomedical research.

The impetus for surveying the graduate training climate of our school originated in fall 2019. Its first objective was to assess mentoring. The goal expanded in early 2020 to include questions about career planning, wellness, resilience, and overall program satisfaction. Development of the survey was initiated by the Associate Dean for Graduate Studies, with input from the entire graduate office staff and from the School of Medicine’s Graduate Council. The council includes the graduate program directors and the president of the Biomedical Graduate Student Association. In early March, the authors of the present report applied for pilot funding to support a new program to enhance career preparation by late stage graduate students. The project was designed to begin with baseline surveys of students and faculty. Two weeks after submitting the proposal, the Covid-19 pandemic prompted the University to suspend all in-person learning through classes and laboratory research. After the pilot grant was approved in June 2020, additional survey elements were added to probe the pandemic’s impact. Before distributing the two surveys, they were submitted to the Institutional Review Board (IRB), which classified them as exempt from further review. The IRB made several suggestions for minor revisions that were incorporated into the final questionnaires. The surveys of students and faculty were distributed on July 9, 2020 and closed on July 31, 2020. During the survey period in July, the University of Pittsburgh maintained a continuing operational posture of ‘elevated risk’ (https://www.coronavirus.pitt.edu/operational-postures). All laboratory research was in early stages of resumption at greatly reduced staffing levels with enhanced safety measures, while summer term classes continued through remote learning over the internet.

### Methodology

The student survey contained 50 items, some of which had multiple components. The faculty survey had 38 items. They were constructed and implemented using Qualtrics^XM^ (Provo, UT) through the University of Pittsburgh web portal. The surveys were sent to all PhD students in School of Medicine programs and to all training faculty affiliated with the programs. Although participants received two email reminders between July 9 and July 31, participation in the survey was voluntary. After giving informed consent, participants answered multiple choice questions and were given access to text boxes where they could make comments to elaborate their answers. Participants were asked to give their names. Students were asked the names of their dissertation advisors. The purpose was to permit longitudinal analysis of individual student trajectories at later time points in their careers and to verify that each respondent was unique. Analysis of the surveys was blinded by assigning a numerical code to each name and then removing the names from the Excel downloads of the data from Qualtrics. One author did the blinding prior to looking at the data, and all three authors analyzed the anonymized data.

## Results

### The Student Survey

The survey was completed by 135 of 350 PhD students, a return rate of 39%. Recipients included all 315 PhD students registered in the School of Medicine and 35 students enrolled in programs jointly administered with other schools. These included 31 students from the Neuroscience and the Molecular Biophysics & Structural Biology joint programs who were registered in the Dietrich School of Arts and Sciences, and 4 Molecular Biophysics & Structural Biology students who were registered at Carnegie Mellon University. Only 10 of the 35 students registered in other schools completed the survey. Thus 7% of the 13respondents were not registered in the School of Medicine.

### Demographics

Of 135 respondents 59% identified as White, 23% as Asian, 11% as Hispanic/LatinX, and 7% as Black. Of these, 7% identified as belonging to multiple races and/or ethnicities. This breakdown resembles the overall demographics of the total of 315 PhD students who were registered in the School of Medicine in July - 52% White, 21% Asian, 8% Hispanic/LatinX, 4% Black, 2% multiple races and/or identities, and 12% unknown.

Of 135 respondents, 62% identified as female and 36% as male. In addition to identifying as either male, female or non-binary, 16% of students identified as LGBTQ+ and/or other.

The academic year was nearing its end when the survey was administered in July. Beginning students were finishing their first year. They had completed laboratory rotations and were transitioning to work under dissertation advisors to start the second year. Rising third year students had in most cases completed their comprehensive examination and were moving on to develop thesis proposals and form dissertation committees. **Early stage PhD trainees**, classified as entering either their second or third years, represented 41% of 134 respondents. **Late stage PhD trainees,** classified as entering their fourth year or beyond, represented 59% of respondents. By comparison, the breakdown of all students registered in the School of Medicine was 49% early stage and 51% late stage.

Students from all 11 PhD-granting programs responded to the survey (Table 1). Variations in participation from different programs reflected program size in most cases. For example, the 7 respondents from Biomedical Informatics represented 5% of survey respondents and 5% of the school’s total enrollment were in this program. For program descriptions see https://somgrad.pitt.edu/.

**Table 1.** PhD Program.

| Table 1. PhD Program | # of survey respondents | % of survey respondents | program as % of school's total enrollment |
| --- | --- | --- | --- |
| Biomedical Informatics | 7 | 5 | 5 |
| Cell Biology and Molecular Physiology | 7 | 5 | 3 |
| Cellular and Molecular Pathology | 14 | 10 | 8 |
| Clinical and Translational Science | 2 | 1 | 5 |
| Computational Biology | 8 | 6 | 10 |
| Integrative Systems Biology | 9 | 7 | 9 |
| Microbiology and Immunology | 30 | 22 | 23 |
| Molecular Biophysics and Structural Biology | 6 | 4 | 3 |
| Molecular Genetics and Developmental Biology | 11 | 8 | 6 |
| Molecular Pharmacology | 17 | 13 | 9 |
| Neurobiology/Neuroscience | 24 | 18 | 19 |

Survey respondents also included 10 MD/PhD students who were registered as graduate students in the School of Medicine. This represents 28% of the 36 MD/PhD students currently enrolled as graduate students in medical school programs.

To obtain a qualitative summary of written comments in this and subsequent sections of the survey, we identified themes in the comments and then counted the number of respondents mentioning each theme. Using this approach, 17 students criticized the wording of the demographic questions regarding race, ethnicity, sex, gender, and sexuality. The most common concerns were that the answer options were too limited and that they conflated distinctions between classifications that should be separate (e.g. sex vs. sexuality). The authors appreciate these concerns and acknowledge the combined categories. The intent of questions with limited options was to obtain classifications that would include the largest number of individuals while avoiding groups of less than 3 people, which might reveal individual identities. This was considered especially important because information about gender identity and sexual preferences is very personal, therefore making it essential to protect individual privacy. Similar logic extended to the question about race and ethnic identity. The purpose of these questions was to learn about the educational climate experienced by different groups. For example, are there racial, ethnic, gender, or sexual identity-based disparities and are such disparities associated with bias, micro-aggressions, and mentoring?

In this and subsequent sections of the written comments, many respondents asked why they needed to disclose their names. There were two reasons. First, to verify that each survey response represented a unique individual. More importantly, the names of trainees were recorded to enable future longitudinal studies of career outcomes. How do student views evolve during graduate school and subsequent entry into the workforce? Consistent with this goal, NIH T32 training grants now require grantees to track outcomes for 15 years after graduation. This survey will become an important piece of career tracking that will enable the school to assess training of all students, not just those supported by training grants. To protect the privacy of individuals, the present analysis was conducted blind. The approach mirrors that used in clinical trials of experimental therapies. In this case the treatment pertains to methods for enhancing the graduate training climate.

When considering data in the following sections, it is important to note that although the demographics of respondents reflect the overall population of students and of faculty in the school, they are not random samples. Instead they are convenience samples because they reflect the views of individuals who chose to complete voluntary surveys. Although this approach limits the use of statistical methods and should be considered exploratory, the sample sizes were sufficient to reveal major trends that warrant more detailed attention.

### Career Goals

Students were asked to prioritize their long-term career goals with questions based on a condensed version of the three-tier taxonomy of career outcomes that was developed by members of the BEST consortium [8] and adopted by the Coalition for Next Generation Life Sciences (https://nglscoalition.org/progress/). The tiers describe “Workforce/Employment Sector”, “Career/Job Type”, and “Job Function”. In our questionnaire we used one tier that contained 5 employment sectors (n=129 respondents) and a second tier with 11 job functions (n= 134 respondents).

Based on student rankings of preferred employment sector, the first choices were academia (36%), business (35%), undecided (22%), government (5%), and other non-profit (3%)

The first choice long-term jobs were tenure track professor at an R1 university (31%), research & development in industry (26%), undecided (14%), other (8%), staff scientist at R1 university (6%), and teaching with a lab at a small college (5%). The other jobs preferred by 1 to 4% of students were intellectual property law, business of science, scientific publishing, teaching professor at small college, science policy, science related sales and customer education.

Students were also asked about their preference for a first position immediately after graduate school. They could check multiple jobs. Table 2 shows the number of times the choices for first position after graduate school were chosen by 135 respondents. The most frequent choice was postdoctoral research training with 42% of the responses.

**Table 2.** preferred first position after graduate school.

| Table 2 – preferred first position after graduate school | # of total responses | % of total responses |
| --- | --- | --- |
| postdoctoral research training | 96 | 42 |
| entry level job in big pharma or a biotech company | 62 | 27 |
| entry level job in publishing | 7 | 3 |
| teaching job | 12 | 5 |
| policy internship in government or a non-profit organization | 13 | 6 |
| business or law school | 4 | 2 |
| undecided at this time | 28 | 12 |
| blank | 4 | 2 |

To see whether goals change during graduate school, we then broke down the data from Table 2 into responses by early stage and late stage trainees (Table 3). The percentage of total responses for each group revealed that late stage students expressed a reduced interest in postdoctoral training and greater interest in other options.

**Table 3.** preferred first position after graduate school (Early vs Late stage trainees)

| **Table 3 – preferred first position after graduate school**<br>**(Early vs Late stage trainees)** | **% of responses by**<br>**Early Stage trainees** | **% of responses by**<br>**Late Stage trainees** |
| --- | --- | --- |
| postdoctoral research training | 83 | 66 |
| entry level job in big pharma or a biotech company | 13 | 21 |
| entry level job in publishing | 0 | 1 |
| teaching job | 0 | 1 |
| policy internship in government or a non-profit organization | 0 | 1 |
| business or law school | 0 | 3 |
| undecided at this time | 4 | 6 |

In text comments on career goals,15 students indicated they were on track and adapting to disruptions caused by the pandemic. Around 20 students expressed serious concern over the delays and how this would impact their productivity and competitiveness for jobs. Around 15 students expressed concern over the potential negative impacts of economic disruption on funding and the job market. In reading through these comments, one cannot escape the impression that students are very worried about how the uncertainty caused by the pandemic will disrupt their lives and plans for the future.

Multiple-choice questions in the survey’s career goals section asked students about their planning efforts and barriers to planning.

When asked about the time they spend thinking about their plans for careers after graduate school,

- 47% of students think about their futures all the time or a good deal of the time.
- 38% spend a moderate amount of time on it.
- 15% spend minimal or no time on this type of planning.

In a question that permitted multiple responses, students were asked about their perceptions of barriers to beginning the process of career exploration and planning.

- 76% indicated that the task either overwhelms them with too many possibilities or that it takes too much time away from their focus on graduate school, or both.
- On the positive side, 22% reported that their plans were on track and their advisor was supportive.
- 7% reported that their advisor tells them they need to spend more time in the lab and less on diversions.
- 5% reported that their advisors were unsupportive of a career outside of academia.

When asked whether they would be interested in a part-time externship or a brief full-time internship to explore a potential career option,

- 54% of students indicated they were very interested or extremely interested in such opportunities
- 5% indicated that had already started the process.
- 22% expressed moderate interest
- 19% expressed little or no interest

In summer 2019, the graduate office hired Dr. McCord to serve as a career exploration and planning specialist. Although some students indicated they were unaware of this change, others were aware and in written comments they expressed enthusiasm for her efforts to date. Further efforts to develop career planning services for PhD students in the medical school therefore appear as an important area for future dialog and action.

### Wellness and Resilience

Graduate school and careers in biomedical science are inherently stressful [9]. They involve continual competition, failure and rejection, punctuated by periodic successes whose rewards can make it all worthwhile. Persisting in science and science-based careers requires that one develop habits and skills that promote wellness and resilience. Working through the problems inherent in one’s work and one’s outside life can lead in the long-term to feelings of accomplishment and satisfaction. Questions in this survey section were designed to probe student work habits and ask how they were dealing with the stresses and strains that are normal parts of life in graduate school and beyond. A total of 132 students completed this section of the survey. Responses considered positive climate indicators are highlighted in green, negative indicators are highlighted in red. The actual language of survey questions is italicized.

Although 95% of the graduate students find time for recreation and other forms of self-care, 37% indicate it is insufficient.

*Are you able to find time for activities outside your studies and research? (examples include recreation, exercise, hobbies, social life, movies, music, theater, cooking)*

| Response | # of responses | % of responses |
| --- | --- | --- |
| No | 6 | 5 |
| a little, but not enough | 49 | 37 |
| Yes | 77 | 58 |

Graduate students work long hours. Before the pandemic 58% worked 50 hours or more per week. Working more than 60 hours per week carries an increased risk of being detrimental to wellness and should be further examined when pandemic restrictions end.

*How many hours per week do spend either in the lab or on other campus activities related to your studies?*

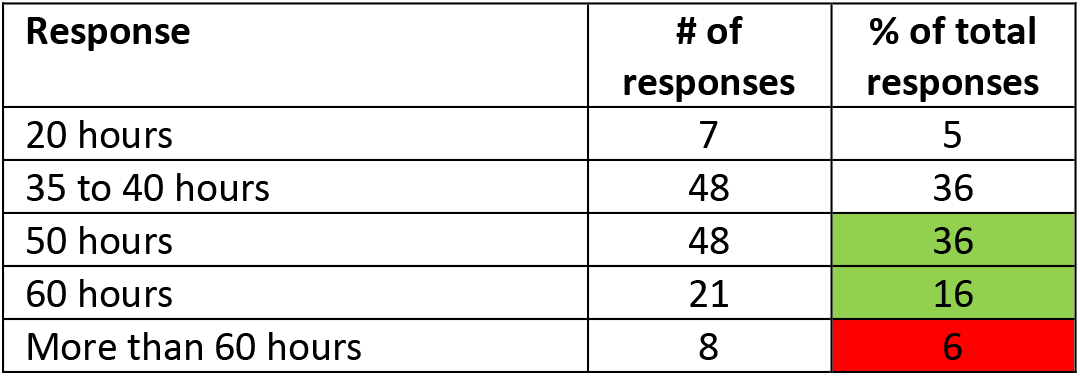

The pandemic has negatively impacted on hours worked by graduate students.

*During the COVID-19 pandemic lock-down, how many hours per week have you spent working remotely from home?*

The pandemic has negatively impacted on hours worked by graduate students. *During the COVID-19 pandemic lock-down, how many hours per week have you spent working remotely from home?*
| Response | # of responses | % of total responses |
| --- | --- | --- |
| 20 hours | 37 | 28 |
| 35 to 45 hours | 71 | 54 |
| 50 hours | 15 | 12 |
| 60 hours | 4 | 3 |
| More than 60 hours | 4 | 3 |

Differences in hours worked in lab before and after 4 months of the COVID lockdown reveal a reduction in research activity by graduate students

| Response | Change in # of students |
| --- | --- |
| 20 hours | Up 30 |
| 35 to 45 hours | Up 23 |
| 50 hours | Down 33 |
| 60 hours | Down 17 |
| More than 60 hours | Down 4 |

Most graduate students often work nights and weekends in their labs.

*How often do you work in the lab on weekends? (before the Pandemic)*

Most graduate students often work nights and weekends in their labs. *How often do you work in the lab on weekends? (before the Pandemic)*
| Response | # of responses | % of total responses |
| --- | --- | --- |
| Never | 11 | 8 |
| Sometimes | 55 | 42 |
| Half the time | 33 | 25 |
| Most of the time | 24 | 18 |
| Always | 9 | 7 |

*How often do you arrive at the lab before 8 AM or leave after 6 PM? (before the Pandemic)*

50% *How often do you arrive at the lab before 8 AM or leave after 6 PM? (before the Pandemic)*
| Response | # of responses | % of total responses |
| --- | --- | --- |
| Never | 5 | 4 |
| Sometimes | 56 | 46 |
| Half the time | 30 | 25 |
| Most of the time | 32 | 26 |
| Always | 9 | 7 |

50% of students experience the imposter syndrome on a regular basis!!

*Sometimes students say, “I feel like an impostor. Everyone else is smarter and I was admitted to the program by mistake.” Do you ever feel this way?*

| Response | # of responses | % of total responses |
| --- | --- | --- |
| Never | 17 | 13 |
| Sometimes | 49 | 37 |
| Half the time | 20 | 15 |
| Most of the time | 27 | 21 |
| Always | 19 | 14 |

The survey included the GAD-2 and PHQ-2 questions, which are used clinically to screen for anxiety and depression [10, 11]. Answers from PhD students covered the full range of possible responses.

*Over the last 2 weeks, how often have you been bothered by the following problems?*

| Response (n=132) | Feeling nervous, anxious or on edge |  | Not able to stop or control worrying |  | Little interest or pleasure in doing things |  | Feeling down, depressed, or hopeless |  |
| --- | --- | --- | --- | --- | --- | --- | --- | --- |
|  | # | % | # | % | # | % | # | % |
| Not at all | 14 | 11 | 38 | 29 | 50 | 38 | 54 | 41 |
| Several Days | 54 | 41 | 51 | 39 | 51 | 39 | 50 | 38 |
| More than half the days | 34 | 26 | 23 | 17 | 22 | 17 | 20 | 15 |
| Nearly every day | 30 | 23 | 20 | 15 | 9 | 7 | 8 | 6 |

Interpretation of the responses requires that each question be scored on a scale of 0-3 and that the scores for each person be added for each pair of questions, yielding numbers from 0-6. Table 4 shows the distribution of individual scores for the two tests.

**Table 4.**

| **Table 4**<br>**Score:** | **0** | **1** | **2** | **3** | **4** | **5** | **6** |
| --- | --- | --- | --- | --- | --- | --- | --- |
| **GAD-2**<br>**anxiety** | 9 | 27 | 35 | 15 | 20 | 9 | 17 |
| **PHQ-2**<br>**depression** | 41 | 19 | 33 | 19 | 12 | 2 | 6 |

Scores ≥3 on either test indicate a likelihood of clinically significant anxiety or depression that may require further evaluation. Table 5 shows the number of graduate students who had a score ≥3 on each test, and on both. The data reveal that **51% of students scored ≥3 on either one or both tests.**

**Table 5.**
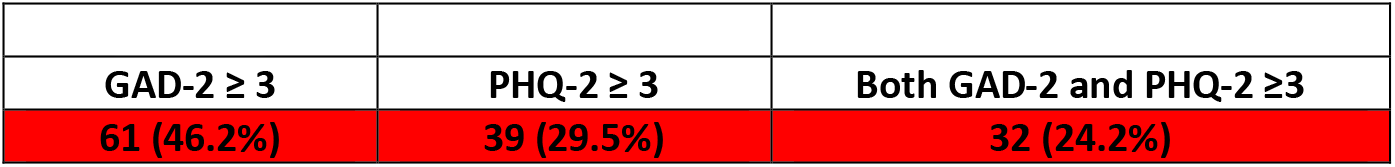

| **Table 5** |  |  |
| --- | --- | --- |
| **GAD-2 $\geq 3$** | **PHQ-2 $\geq 3$** | **Both GAD-2 and PHQ-2 $\geq 3$** |
| **61 (46.2%)** | **39 (29.5%)** | **32 (24.2%)** |

A much smaller proportion of graduate students appear to utilize mental health services through the University counseling center. Only 24 students answered this question. Their written comments indicate they are not satisfied with the counseling center.

*Over the past year, the University has invested in strengthening the counseling center. Have you ever used any of the services they offer? Check all that apply.*

| **Response** | **# of responses** | **% of respondents (n= 24)** | **% total group completing the survey (n= 132)** |
| --- | --- | --- | --- |
| Drop-in hours during the week | 12 | 50 | 9 |
| Group sessions | 0 | 0 | 0 |
| Focused workshops | 1 | 4 | 1 |
| Referrals to see a physician | 2 | 8 | 2 |
| No. I never use these services | 0 | 0 | 0 |
| **Combinations:** |  |  |  |
| Drop-in + group sessions | 6 | 25 | 5 |
| Drop-in + physician referrals | 1 | 4 | 1 |
| Group + physician referrals | 1 | 4 | 1 |
| Drop in + group + referral | 1 | 4 | 1 |

The School of Medicine has taken steps to establish an Ombuds office, with capacity to help graduate students. 37% of the graduate students are interested in this support mechanism.

*How likely is it that you may want to speak with the SOM Ombudsperson?*

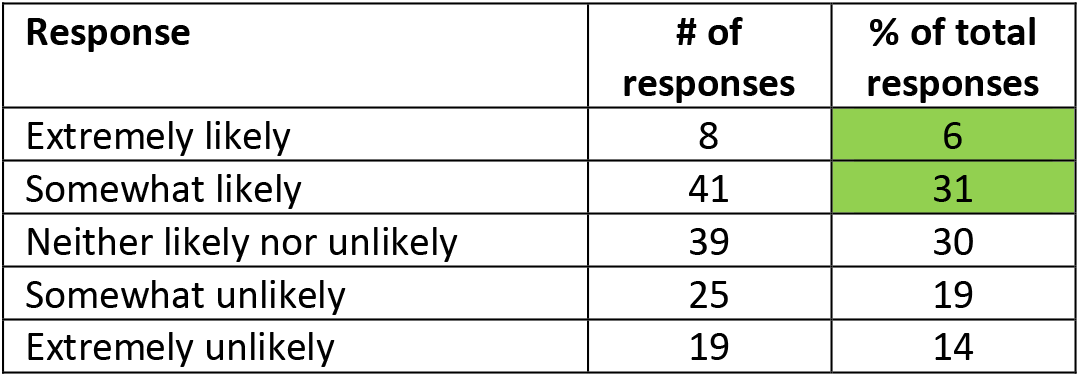

| **Response** | **# of responses** | **% of total responses** |
| --- | --- | --- |
| Extremely likely | 8 | 6 |
| Somewhat likely | 41 | 31 |
| Neither likely nor unlikely | 39 | 30 |
| Somewhat unlikely | 25 | 19 |
| Extremely unlikely | 19 | 14 |

Responses to several questions revealed problems with bias. 47% of medical school graduate students believe we could do a much better job in embracing diversity and inclusion!

*To what extent do you believe that graduate programs in the School of Medicine have created a culture that embraces diversity and inclusion?*

| **Response** | **# of responses** | **% of total responses** |
| --- | --- | --- |
| Not at all | 13 | 10 |
| A little | 49 | 37 |
| Sometimes | 46 | 35 |
| Definitely yes | 24 | 18 |

30% of graduate students do not feel welcome in our community.

*“People like me feel like they are welcome as members of the medical school community”*

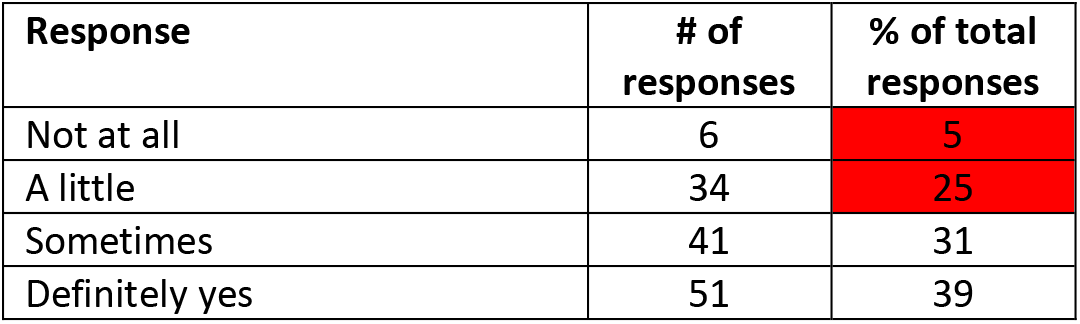

45% of graduate students have never had implicit bias training.

*How many times have you participated in a workshop or online training on the topic of implicit bias?*

45% of graduate students have never had implicit bias training. *How many times have you participated in a workshop or online training on the topic of implicit bias?*
| Response | # of responses | % of total responses |
| --- | --- | --- |
| Never | 59 | 45 |
| Once | 48 | 36 |
| Twice | 16 | 12 |
| Three or more times | 9 | 7 |

24% of graduate students have been subject to microaggressions

*How often have you been exposed to microaggressions in the context of your graduate education at Pitt? - Either on the receiving end or as a witness?*

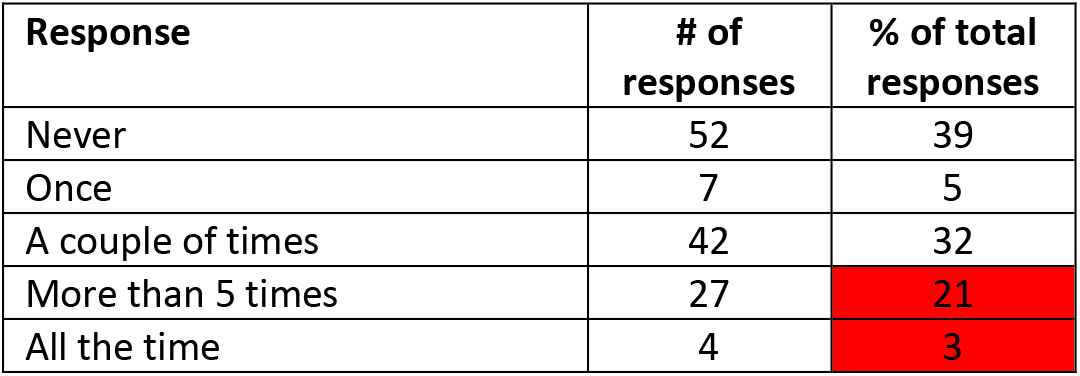

| Response | # of responses | % of total responses |
| --- | --- | --- |
| Never | 52 | 39 |
| Once | 7 | 5 |
| A couple of times | 42 | 32 |
| More than 5 times | 27 | 21 |
| All the time | 4 | 3 |

### Mentoring

Building strong professional relationships between PhD students and their mentors is believed to be a powerful driver of academic success, research productivity, and satisfaction. This section of the survey probed these relationships.

72% of PhD students reported having at least one secondary mentor

*In addition to their dissertation adviser, students sometimes develop mentoring relationships with other faculty. Sometimes these secondary mentors are from one’s thesis committee. Other times they are from the student’s program, a journal club or are lab neighbors. How many secondary mentors do you have besides your dissertation adviser?*

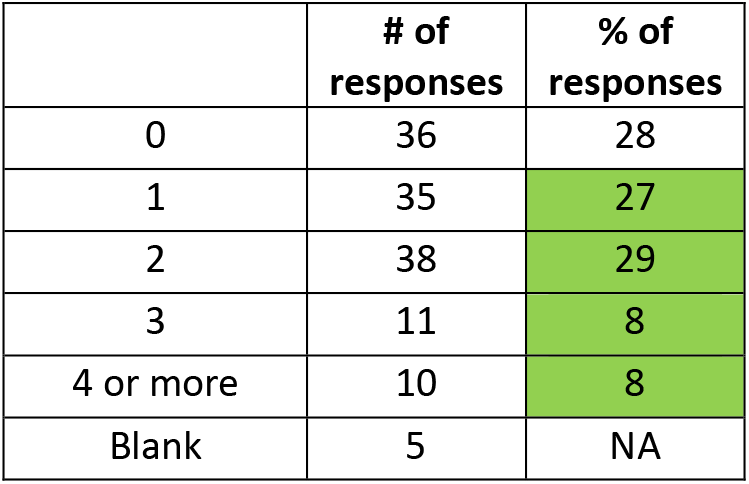

After removing second year students who are just joining dissertation labs from the analysis, 76% of the more experienced students reported having more than one secondary mentor

|  | # of responses | % of responses |
| --- | --- | --- |
| 0 | 22 | 22 |
| 1 | 25 | 26 |
| 2 | 31 | 32 |
| 3 | 11 | 11 |
| 4 or more | 9 | 9 |
| Blank | 3 | NA |

58% of students report meeting individually with their dissertation advisors ≥ once per week. On the negative side, 19% report seeing their mentor once a month or less.

*On average, my dissertation adviser meets individually with me for 15 minutes or more*

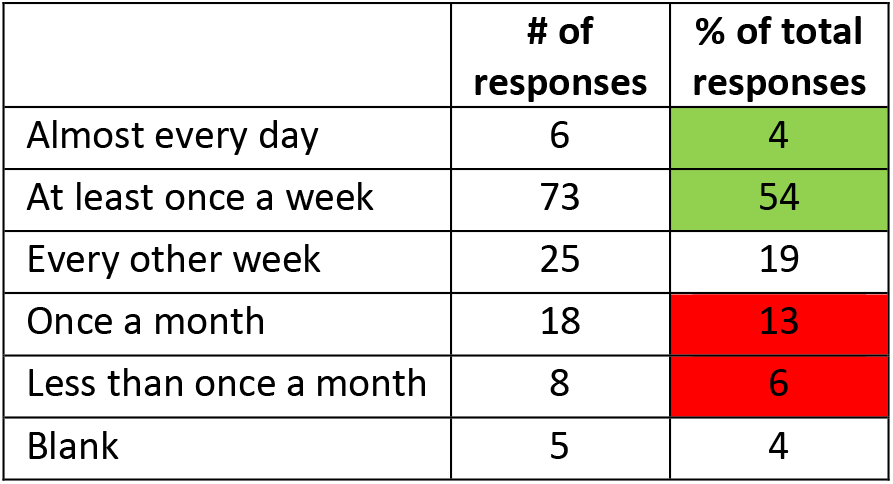

|  | # of responses | % of total responses |
| --- | --- | --- |
| Almost every day | 6 | 4 |
| At least once a week | 73 | 54 |
| Every other week | 25 | 19 |
| Once a month | 18 | 13 |
| Less than once a month | 8 | 6 |
| Blank | 5 | 4 |

89% of students never use the AAMC mentoring compact, even though it is distributed to most of them.

*The AAMC (American Association of Medical Colleges) publishes a compact to guide discussions between graduate students and their mentors. How many times have you used this compact in discussions with your dissertation adviser?*

|  | # of responses | % of total responses |
| --- | --- | --- |
| never | 120 | 89 |
| once | 9 | 7 |
| multiple times | 1 | 1 |
| Blank | 5 | 4 |

Most students (70 to 88%) gave high marks when rating their dissertation advisors on an 11-point scale. For simplicity, the scores were binned into three groups (red, orange, green), representing low to high marks.

*My dissertation adviser cares about me as a person.*

*My dissertation adviser treats my scientific ideas and insights with respect.*

*My adviser gives me freedom to chart the direction of my dissertation research.*

*My dissertation adviser and I are on the same page when it comes to understanding each other’s mutual expectations.*

*When I have a problem, either personal or professional, my dissertation advisor displays empathy. Even if they cannot solve the problem, they are willing to listen.*

**SCALE: (totally true = 10, totally false = 0)**

|  | Cares about me as a person |  | Respects my scientific ideas |  | Freedom to chart research direction |  | Same page on mutual expectations |  | Displays empathy |  |
| --- | --- | --- | --- | --- | --- | --- | --- | --- | --- | --- |
|  | # of responses | % of responses | # of responses | % of responses | # of responses | % of responses | # of responses | % of responses | # of responses | % of responses |
| 0 | 3 | 2 | 0 | 0 | 0 | 0 | 1 | 1 | 1 | 1 |
| 1 | 0 | 0 | 0 | 0 | 3 | 2 | 2 | 2 | 1 | 1 |
| 2 | 3 | 2 | 3 | 2 | 1 | 1 | 2 | 2 | 2 | 2 |
| 3 | 1 | 1 | 4 | 3 | 2 | 2 | 5 | 4 | 4 | 3 |
| 4 | 0 | 0 | 2 | 2 | 3 | 2 | 5 | 4 | 4 | 3 |
| 5 | 6 | 5 | 9 | 7 | 17 | 13 | 10 | 8 | 10 | 8 |
| 6 | 3 | 2 | 3 | 2 | 10 | 8 | 13 | 10 | 7 | 6 |
| 7 | 11 | 9 | 15 | 12 | 17 | 13 | 21 | 16 | 12 | 9 |
| 8 | 17 | 13 | 16 | 12 | 18 | 14 | 19 | 15 | 14 | 11 |
| 9 | 25 | 19 | 20 | 16 | 20 | 16 | 24 | 19 | 16 | 13 |
| 10 | 61 | 47 | 57 | 44 | 38 | 30 | 26 | 20 | 57 | 45 |
| **Sum of 7-10** |  | **88%** |  | **84%** |  | **73%** |  | **70%** |  | **78%** |

9% of students report being criticized in front of others using counterproductive methods. 86% are not treated this way.

*My dissertation adviser has criticized my work in front of others in a way that I find counterproductive*

9% of students report being criticized in front of others using counterproductive methods. 86% are not treated this way. *My dissertation adviser has criticized my work in front of others in a way that I find counterproductive*
| Frequency of behavior | # of responses | % of total responses |
| --- | --- | --- |
| never | 99 | 73 |
| once | 18 | 13 |
| repeatedly (more than 3 times) | 12 | 9 |
| blank | 6 | 4 |

87% of mentors are <u>not</u> screamers, but some engage in this negative practice.

*My dissertation adviser gets angry, raises their voice and yells at people*

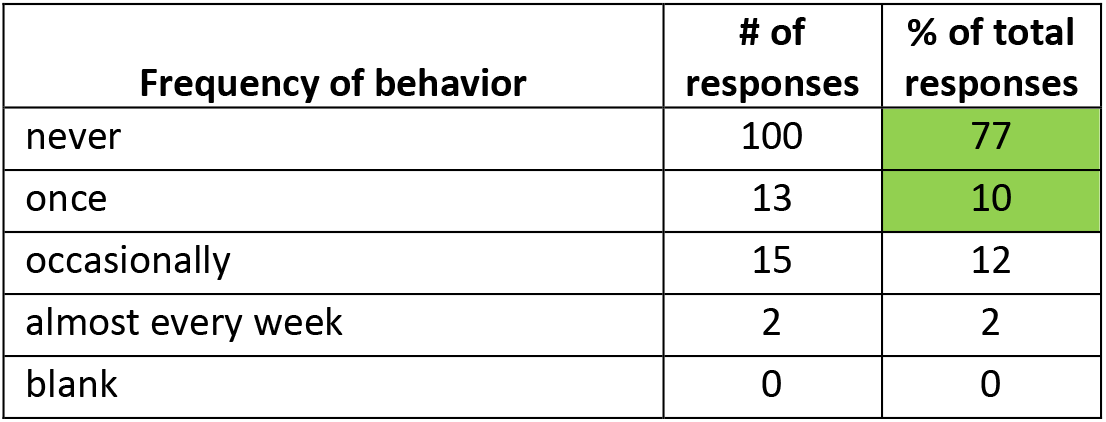

87% of mentors are not screamers, but some engage in this negative practice. *My dissertation adviser gets angry, raises their voice and yells at people*
| Frequency of behavior | # of responses | % of total responses |
| --- | --- | --- |
| never | 100 | 77 |
| once | 13 | 10 |
| occasionally | 15 | 12 |
| almost every week | 2 | 2 |
| blank | 0 | 0 |

Most, but not all, graduate students attend scientific meetings. The late stage students (years 3 and beyond) who never or rarely attend conferences indicate a problem.

*How many national scientific meetings have you attended and presented an abstract*

| # of meetings | # of responses | % of total responses |
| --- | --- | --- |
| none | 37 | 28 |
| 1 | 31 | 24 |
| 2 | 30 | 23 |
| 3 | 10 | 8 |
| 4 or more | 22 | 17 |

| # of meetings | second year | third year | fourth year | fifth year | sixth year or beyond |
| --- | --- | --- | --- | --- | --- |
| none | 22 | 9 | 3 | 3 | 0 |
| 1 | 5 | 6 | 6 | 8 | 6 |
| 2 | 2 | 3 | 11 | 9 | 5 |
| 3 | 2 | 1 | 2 | 3 | 2 |
| 4 or more | 1 | 2 | 7 | 7 | 5 |

55% of mentors consistently help their graduate students attend meetings, but 45% eschew this important practice. The problem of not supporting students to attend meetings extends into latter years of training.

*My dissertation adviser has encouraged and facilitated my participation in national scientific meetings*

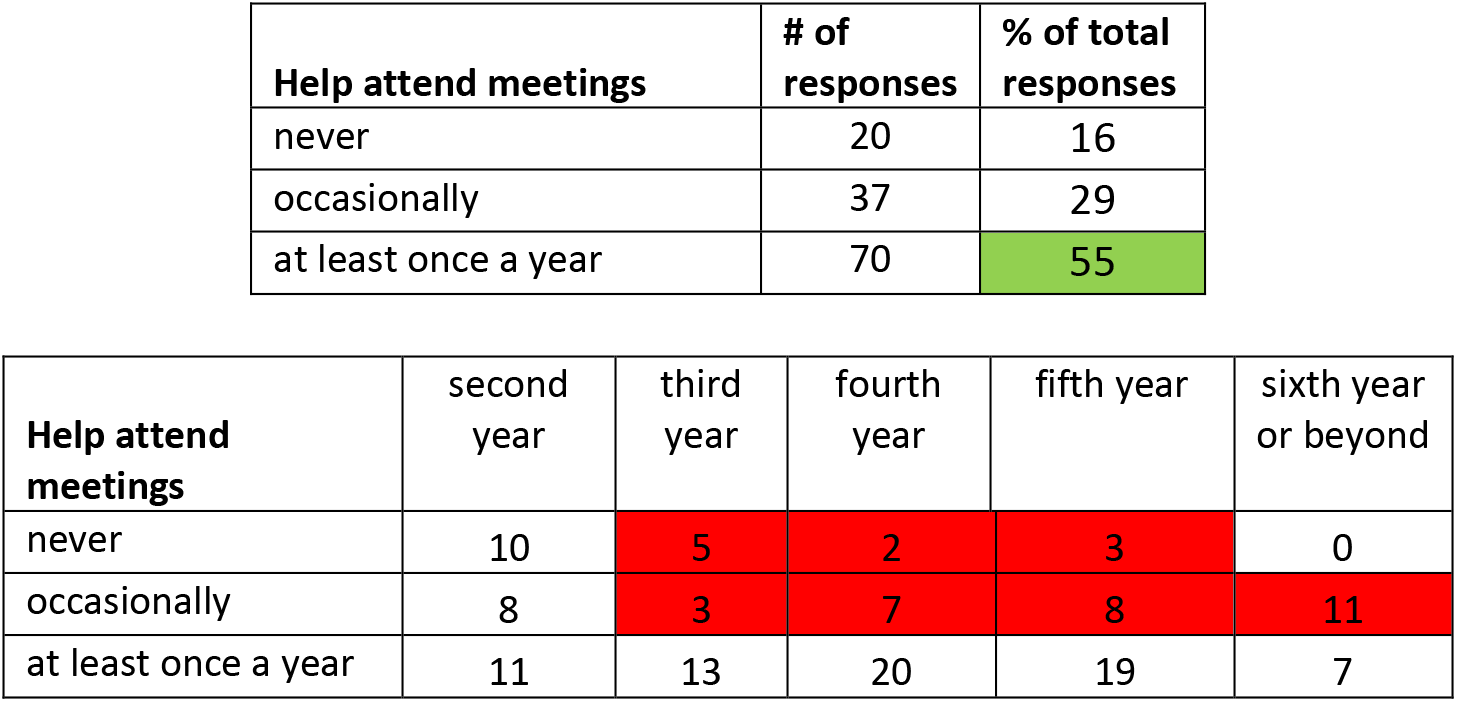

*63% of mentors <u>rarely</u> introduce their students to established scientists!*

*At scientific meetings, my dissertation adviser has introduced me to established scientists in my field who work at other universities.*

| Introduce students to established scientists | # of responses | % of total responses |
| --- | --- | --- |
| never | 49 | 40 |
| occasionally | 36 | 29 |
| at every meeting | 38 | 31 |

The lack of professional socialization extends throughout all stages of training.

| Introduce students to established scientists | second year | third year | fourth year | fifth year | sixth year or beyond |
| --- | --- | --- | --- | --- | --- |
| never | 19 | 7 | 9 | 9 | 5 |
| occasionally | 6 | 6 | 6 | 10 | 8 |
| at every meeting | 3 | 5 | 14 | 11 | 5 |

71% of mentors help their students develop writing skills and 68% are familiar enough with program guidelines to help their students stay on track.

*My dissertation adviser has spent time helping me become a better scientific writer*

*My dissertation adviser is familiar with the guidelines and milestones of my program and helps me stay on track.*

**Score: totally true = 10, totally false = 0**

| score | Writing |  | Program guidelines |  |
| --- | --- | --- | --- | --- |
|  | # | % | # | % |
| 0 | 1 | 1 | 0 | 0 |
| 1 | 1 | 1 | 4 | 3 |
| 2 | 2 | 2 | 4 | 3 |
| 3 | 1 | 1 | 5 | 4 |
| 4 | 0 | 0 | 4 | 3 |
| 5 | 13 | 10 | 8 | 6 |
| 6 | 12 | 10 | 10 | 8 |
| 7 | 21 | 17 | 14 | 11 |
| 8 | 24 | 19 | 22 | 17 |
| 9 | 10 | 8 | 16 | 13 |
| 10 | 41 | 33 | 39 | 31 |

### Overall Satisfaction

Based on simple check boxes, large proportions of the students report being moderately or extremely happy with their PhD program (65%), their dissertation advisor (84%), their dissertation project (74%), and their department or center (73%). However, only 39% of students are extremely or moderately happy about their feelings of belonging to the medical school community and 58% to the University at large. Note that 31% of graduate students were unhappy about their feelings of inclusion in the medical school community

*Putting it all together, I am happy with these elements of my training:*

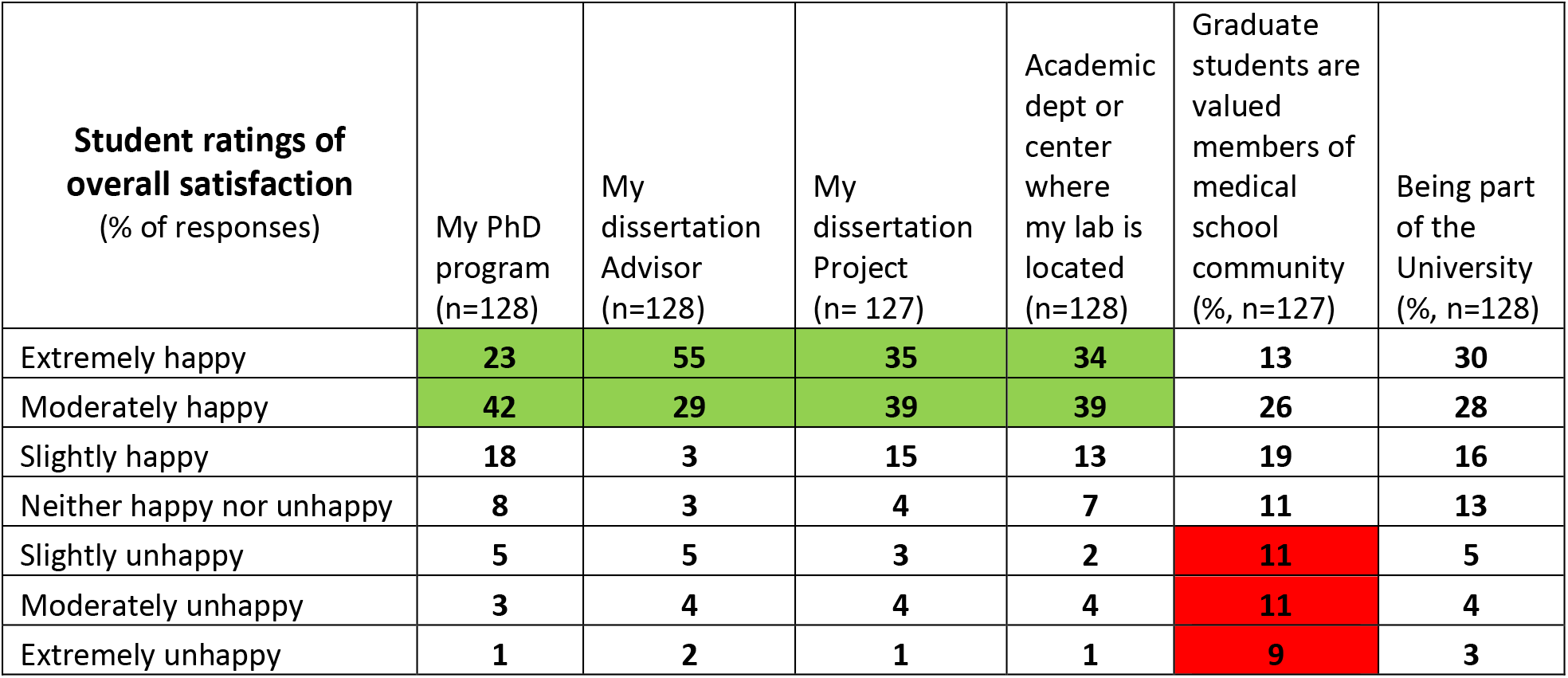

| Student ratings of overall satisfaction<br>(% of responses) | My PhD program<br>(n=128) | My dissertation Advisor<br>(n=128) | My dissertation Project<br>(n=127) | Academic dept or center where my lab is located<br>(n=128) | Graduate students are valued members of medical school community<br>(%, n=127) | Being part of the University<br>(%, n=128) |
| --- | --- | --- | --- | --- | --- | --- |
| Extremely happy | 23 | 55 | 35 | 34 | 13 | 30 |
| Moderately happy | 42 | 29 | 39 | 39 | 26 | 28 |
| Slightly happy | 18 | 3 | 15 | 13 | 19 | 16 |
| Neither happy nor unhappy | 8 | 3 | 4 | 7 | 11 | 13 |
| Slightly unhappy | 5 | 5 | 3 | 2 | 11 | 5 |
| Moderately unhappy | 3 | 4 | 4 | 4 | 11 | 4 |
| Extremely unhappy | 1 | 2 | 1 | 1 | 9 | 3 |

### The Faculty Survey

The survey was completed by 83 of 414 members of the School of Medicine graduate program faculty, a return rate of 20%.

### Demographics

Of the 83 respondents, 78% identified as White, 14% as Asian, 5% as Hispanic/LatinX, and 2% as Black/African American. Additionally, 30% identified as female, 69% as male, and 1% as LGBTQ+ and/or other. The majority (70%) of faculty respondents have been at Pitt under 15 years. Faculty associated with all School of Medicine Graduate Training Programs responded. Note, some faculty are associated with multiple graduate programs, and some faculty did not respond to this question. About half the faculty respondents (51%) were affiliated with 1 SOM graduate program, and the other half with 2 or more SOM graduate programs.

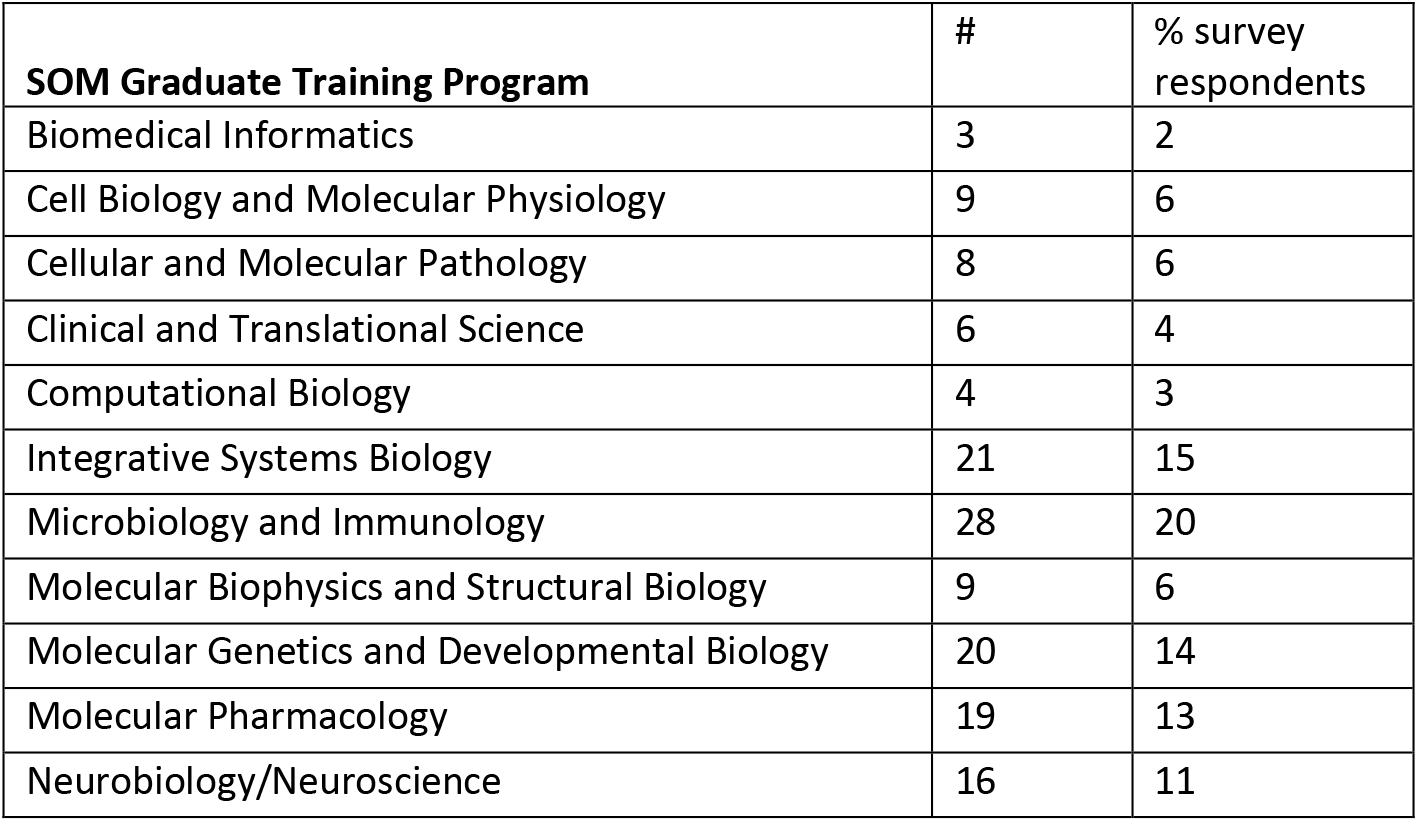

### Career Exploration and Planning

Based on responses to a multiple-choice question, 46% of faculty believe their students spend either a great deal or a lot of time thinking about their career plans. While this corresponds to the student’s responses (47%) to the same question, it is not directly comparable, since the populations are not matched. In written comments, 83% of faculty indicated strong support for their students to attend professional development activities. Their suggestions centered around career workshops/seminars, development of grant writing skills, networking, and technical or analytical skills. They also expressed support of skill building for teaching, public speaking, and outreach. Although many faculty indicated support for these forms of career development, they indicated that they wait until students approach them about the directions they want to take for career exploration. This suggests that future efforts might stress helping faculty to take a more pro-active stance on career development issues.

51% the faculty respondents expressed strong support for student externships and internships. This echoed student responses, where 54% indicated interest in externships and internships

*How strongly would you encourage graduate students who have advanced to candidacy to participate in a part-time externship or a brief full-time internship in order to explore a potential career option?*

|  | # (n= 80) | % |
| --- | --- | --- |
| I would discourage this. Students need to focus on their dissertation research | 7 | 9 |
| I do not have a strong opinion on this question and would wait for a student to raise the issue. | 20 | 25 |
| I would consider but have reservations about disruption of my R01-funded research program | 12 | 15 |
| I would enthusiastically support, as long as we could coordinate our schedules and goals | 41 | 51 |

In written comments, many of the faculty voiced concerns about the effect of the pandemic on the training timelines and careers of their students, including the impact it has had on trainee stress-levels.

### Wellness and Resilience

Only 56% of faculty respondents were aware of resources available to our trainees via the University’s counseling center, or the effort to appoint an Ombudsperson for graduate students. 25% of faculty believe we can do better in embracing diversity and inclusion (compared to the 46% of students who feel similarly). Only 12% of faculty do not feel welcome in our community, compared to 30% of students. This difference may reflect the greater diversity of the student population. 19% of our faculty have never had implicit bias training, and 31% have had training only once. In written comments, most faculty expressed strong feelings about the need to be more intentional about future efforts to improve diversity and inclusion in our community.

### Mentoring

90% of faculty encourage their students to work with secondary mentors. Again, this echoed strong support by students for engaging with secondary mentors. 87% of the faculty report meeting individually with their students for 15 or more minutes at least once per week. However, 10% report meeting only twice a month with their trainees and 4% report meeting only once a month. This suggests that the regularity of one-on-one time between dissertation advisors and their students merits further scrutiny.

Although 59% of faculty members have attended two or more mentor training workshops, 22% have attended only one such workshop and 20% reported no mentor training.

88% of faculty members reported never using the AAMC compact between graduate students and mentors. On a positive note, 85% believe they have a good understanding of mutual expectations with their students. 91% of faculty respondents reported that they take their students to national meetings at least once a year, and 90% report that they introduce students to established scientists at every meeting. These values were higher than reported by students and should be examined further. 87% of faculty mentors reported they help improve their student’s writing skills, and 92% reported being aware of their graduate program milestones for their students.

### Overall Satisfaction

Faculty were asked to rate the quality of medical school graduate students and the quality of the training faculty. Both scored high. 78% of faculty ranked Pitt SOM graduate students high when compared to other leading research universities. Similarly, quality of PhD training faculty at Pitt SOM was ranked high by 84% of respondents. 69% of faculty responded that during their tenure at Pitt SOM the graduate programs have improved. As to program size, 51% responded the student enrollment at Pitt SOM is about the right size, 33% that it is much too small or a bit small, and 16% that it is a bit large or much too large. Classroom facilities were rated as either grossly inadequate or in need of updates by 43% of the faculty. Another 39% responded that classrooms were adequate and only 19% that these facilities are really good.

Putting it all together in another question, 91% of faculty rank the PhD program structure favorably (over 60 on a scale of 0-100, see below), and 72% feel positive about administrative support provided by the Office of Graduate Studies. 81% are positive about the opportunities for students to develop transferrable skills, but only 60% are positive about opportunities for students to explore career options.

Importantly, only 55% of faculty respondents feel that there is evidence that the PhD students are valued members of the medical school community.

| Overall ratings by faculty (% of respondents) | PhD Program Structure | Support provided by the Office of Graduate Studies | Transferrable skills development for students | Career Exploration opportunities for students | Evidence that PhD students are valued members of the medical school community |
| --- | --- | --- | --- | --- | --- |
| 0-9 | 0 | 0 | 1 | 1 | 4 |
| 10-19 | 1 | 0 | 0 | 1 | 5 |
| 20-29 | 1 | 6 | 1 | 3 | 9 |
| 30-39 | 2 | 1 | 4 | 5 | 1 |
| 40-49 | 1 | 6 | 4 | 5 | 10 |
| 50-59 | 4 | 15 | 10 | 25 | 16 |
| 60-69 | 10 | 10 | 20 | 20 | 7 |
| 70-79 | 25 | 22 | 30 | 20 | 25 |
| 80-89 | 37 | 25 | 17 | 15 | 12 |
| 90-100 | 19 | 15 | 14 | 5 | 11 |

## DISCUSSION

The data reported here reflect the first ever climate assessment of the graduate training environment in our medical school at the University of Pittsburgh. It was conducted at a crucial time in history, four months into major local disruptions caused by the global Covid-19 pandemic. Almost certainly the perceptions of the students and faculty members alike were influenced by the pandemic. The challenge in understanding our educational environment was further compounded by a national awakening in the wake of several high-profile killings of black American citizens by police officers, including the 2018 death of Mr. Antwon Rose in Pittsburgh. All these issues came together, just as our institution underwent a major leadership transition with the onboarding in June 2020 of Dr. Anantha Shekhar, the new Dean of Medicine and Senior Vice Chancellor for Health Sciences.

In this report we summarize key features of the survey data. Interpretative comments in the narrative have by design been kept to a minimum. In our view the next step in building a roadmap for future educational efforts should begin with discussions between the major stakeholders: our students, faculty, and leadership. We see opportunities for change guided by data-driven discussions that build upon on ideas expressed by a large sample of our community. We are optimistic that current conditions have created a fortuitous moment that makes possible significant changes in institutional culture and structure.

With these goals in mind, the present report should be viewed as a first edition of the survey results. We anticipate that discussions within our community will drive secondary analyses of the data. Ideally, such discussions should also serve to help prioritize reforms in how we prepare PhDs for the world they will enter after graduate school.

To initiate the further discussion of data, we begin by noting that some of the results confirm ideas we suspected to be true, but for which there was no data. Here are four brief examples.

1. Knowing that half of our students are experiencing symptoms of anxiety and depression is alarming, but not new. Even before the pandemic, evidence from other convenience surveys indicated that anxiety and depression symptoms are expressed at similarly high rates by graduate students [9] at other universities. Why is it that rates of anxiety and depression symptoms are 5 to 10 times higher amongst graduate students than in age-matched non-student populations? How can we better understand this problem and develop effective mitigation strategies?
2. We also know from earlier tracking of training outcomes in our school, from the 2012 NIH workforce report [3] and from work through the NIH BEST program [1] that many biomedical PhD trainees have been pursuing careers outside academia. Now we have local data to see how these trends are shaping up in 2020. The climate survey data may help frame future efforts to facilitate career planning and exploration by our students, even in the face of massive current uncertainty.
3. Before the survey, we worried that many of our faculty were resistant to mentor training and the added effort required to foster career development by students. Now we have evidence that faculty are open to such change. This suggests that our institutional culture is already changing from what it was ten or twenty years ago. The data also help paint a clearer picture of the rate limiting barriers that must be overcome to better support students and their mentors.
4. Although biomedical research is highly prized, graduate education in academic medical centers, here and elsewhere, generally receives secondary priority. Perhaps it should not be surprising, but the present survey now documents that such secondary status is keenly felt by our graduate students. Many graduate students come to our school motivated by the desire to impact on health care and attracted by the tight integration in our school between fundamental biomedical research and clinical programs. Creative work by graduate students fuels discovery and innovation that helps drive medical progress. Now is an opportune time for reforms that recognize the integral role of graduate students in our academic medical community.

Moving forward, we plan on townhall style meetings with students and faculty to discuss the survey results. We hope that feedback from all stakeholders will give form to future efforts.

## Acknowledgements

This work was supported by a grant from the Burroughs Wellcome Fund and by institutional support from the University of Pittsburgh School of Medicine.

## References

1. Lara, L.I., L. Daniel, and R. Chalkley, eds. BEST 1st Edition: Implementing Career Development Activities for Biomedical Research Trainees. 2020, Academic Press. 282 pages.

2. Lenzi, R.N., et al., The NIH “BEST” programs: Institutional programs, the program evaluation, and early data. FASEB J, 2020. 34(3): p. 3570–3582.

3. Tilghman, S., et al., BIOMEDICAL RESEARCH WORKFORCE WORKING GROUP REPORT. 2012, https://acd.od.nih.gov/documents/reports/Biomedical_research_wgreport.pdf. p. 156 pages.

4. Future of Bioscience Graduate and Postdoctoral Training-White Paper. 2017; Available from: https://gs.ucdenver.edu/fobgapt2/pdf/FOBGAPT2_whitepaper_final.pdf.

5. Hitchcock, P., et al., The future of graduate and postdoctoral training in the biosciences. Elife, 2017. 6.

6. Sorkness, C.A., et al., A new approach to mentoring for research careers: the National Research Mentoring Network. BMC Proc, 2017. 11(Suppl 12): p. 22.

7. Jacob, R.R., et al., The “secret sauce” for a mentored training program: qualitative perspectives of trainees in implementation research for cancer control. BMC Med Educ, 2020. 20(1): p. 237.

8. Stayart, C.A., et al., Applying inter-rater reliability to improve consistency in classifying PhD career outcomes. F1000Res, 2020. 9: p. 8.

9. Evans, T.M., et al., Evidence for a mental health crisis in graduate education. Nat Biotechnol, 2018. 36(3): p. 282–284.

10. Kroenke, K., R.L. Spitzer, and J.B. Williams, The Patient Health Questionnaire-2: validity of a two-item depression screener. Med Care, 2003. 41(11): p. 1284–92.

11. Kroenke, K., et al., The Patient Health Questionnaire Somatic, Anxiety, and Depressive Symptom Scales: a systematic review. Gen Hosp Psychiatry, 2010. 32(4): p. 345–59.

